# A filopodia-based dendritic mechanosensory compartment in CSF-contacting neurons

**DOI:** 10.64898/2026.04.09.713694

**Authors:** Daniel Prieto, María Inés Rehermann, Gabriela Fabbiani, Magdalena Vitar, Omar Trujillo-Cenóz, María Victoria Falco, Mikaela Cúparo, Federico F. Trigo, Raúl E. Russo

## Abstract

Cerebrospinal fluid-contacting neurons (CSF-cNs) are spinal sensory cells that detect chemical and mechanical stimuli via PKD2L1 channels located on a primary cilium, as established in aquatic vertebrates like zebrafish. The mechanosensory mechanism in mammals, however, has remained unclear due to the absence of definitive evidence for cilia on their apical processes (ApPrs). Here we show that mouse CSF-cN ApPrs lack cilia but instead possess drebrin-stabilized filopodia enriched with F-actin. Mechanical stimulation of these cilia-free ApPrs elicits robust, PKD2L1-dependent inward currents, that are sufficient to drive neuronal firing, confirming a novel, cilia-independent mechanosensory mechanism. Comparative analyses indicate an evolutionary divergence from ciliary mechanotransduction, with mice adopting a direct, actin-associated mechanism. These findings advance our understanding of the cellular basis of spinal mechanosensation in mammals and reveal a specialized adaptation for monitoring central canal dynamics, with implications for spinal sensory integration and evolution.

**Graphical Abstract:** 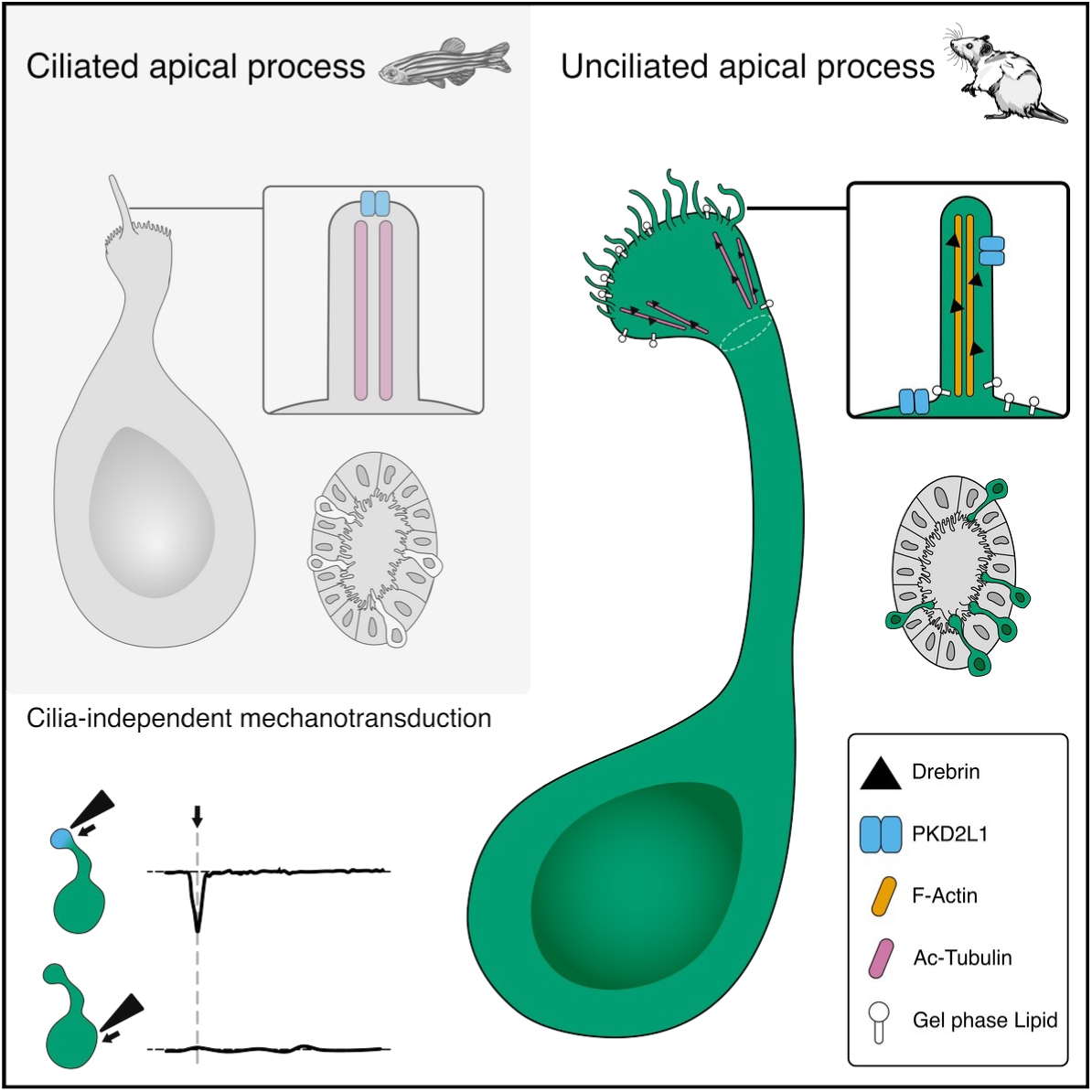

**In Brief:** Cerebrospinal fluid-contacting neurons detect mechanical forces in the spinal cord. In zebrafish, they use a cane-like cilium. Whether mammals inherited this mechanism is controversial. Here, researchers show that mice have replaced the cilium with filopodia, hair-like, touch-sensitive projections stabilized by the protein drebrin. This evolutionary innovation reveals how sensory cells adapt their machinery to the demands of different body plans and environments.

**HIGHLIGHTS:**

- Mouse spinal CSF-cN apical processes lack cilia.
- Drebrin-stabilized filopodia replace the ancestral cilium.
- Direct mechanical stimulation of ApPrs evokes PKD2L1-dependent currents and firing.
- Mammalian CSF-cNs evolved a cilia-independent mechanosensory mechanism.

## INTRODUCTION

Cerebrospinal fluid-contacting neurons (CSF-cNs) are sensory cells located around the central canal (cc) of the vertebrate spinal cord.^1^ They are characterized by a unique anatomy: a single apical dendrite that extends into the cc lumen, terminating in a bulbous apical process (ApPr). This striking structural specialization has long suggested a sensory role, with the ApPr posited as a detector of physicochemical changes within the CSF.^2^

Support for this sensory function comes primarily from studies in non-mammalian vertebrates.^3,4^ In zebrafish, CSF-cN mechanosensation is mediated by the polycystic kidney disease 2-like 1 (PKD2L1) ion channel within a primary cilium projecting from the ApPr,^5,6^ a mechanism essential for controlling spinal curvature during locomotion.^7^ Similar ciliated ApPrs have been described in lamprey and freshwater turtles,^4,8^ establishing a conserved model of ciliary mechanotransduction in aquatic species.

However, extending this ciliary model to mammals is controversial. Although mammalian CSF-cNs express PKD2L1 channels,^9^ consistent with a sensory role, definitive ultrastructural evidence for a motile or primary cilium on their ApPrs is lacking. The ApPr is immersed within a dense meshwork of ependymal cell cilia, obscuring histological interpretation and leading to conflicting reports.^4,9–15^ We therefore asked: Are mammalian CSF-cN ApPrs ciliated? If not, do they retain the mechanotransduction attributed to them?

Here, we clear this discrepancy by demonstrating that mouse CSF-cNs achieve mechanosensation through a novel, cilia-independent mechanism. We show that their ApPrs lack cilia but are instead equipped with drebrin-stabilized filopodia. Mechanical stimulation of ApPrs evokes PKD2L1-dependent currents and the generation of action potentials, confirming the role of the channel in a non-ciliary context. Our findings challenge the current paradigm of CSF-cN mechanoreception and reveal an evolutionary divergence in the receptorial specializations to detect mechanical perturbations within the cord.

## RESULTS

### The apical processes of CSF-cNs lack cilia

Because a ciliary PKD2L1 channel is thought to mediate CSF-cN mechanosensation,^5,15,16^ we first sought to definitively characterize the apically-projecting structures of CSF-cNs in mice. ApPrs extend into the anatomically complex cc lumen where they intermingle with the apical extensions—mainly microvilli and motile cilia—of neighboring ependymal cells. To achieve single cell spatial resolution, we used spinal cord slices from *GATA3^eGFP^*mice,^17^ where we filled either ependymocytes or CSF-cNs with biocytin after patch-clamp recordings to analyze their morphology. As expected from previous work,^14,18^ ependymal cells exhibited 2-4 straight, parallel cilia with no tapering (Figure 1A). The protrusions emanating from GFP^+^ CSF-cN ApPrs were clearly different: they were highly variable in number, length, orientation and morphology with irregular shape (Figure 1B-C).

**Figure 1.**
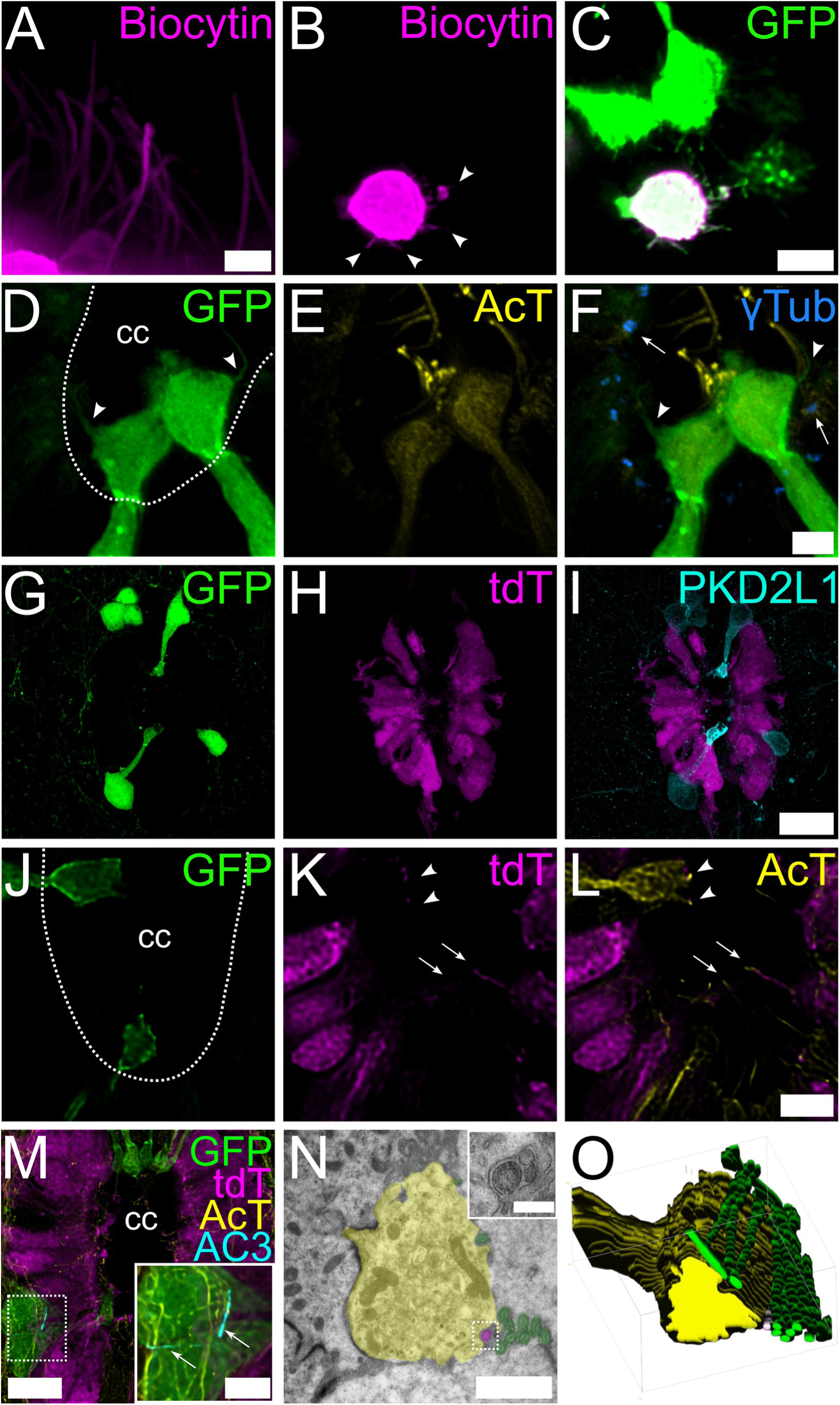
Mouse CSF-cN apical process lack cilia. Morphology of cilia and apical protrusions. (A) A small cluster of biocytin-filled (magenta) ependymocytes with straight, uniform cilia. Maximum intensity projections of deconvolved confocal z-stacks Scale bar, 2µm. (B) A single biocytin-filled CSF-cN from a *GATA3^eGFP^* mouse exhibits numerous irregular protrusions (arrowheads). (C) CSF-cN apical processes (ApPr) protrusions distinct from ependymal cilia. Oblique sections. Scale bar, 5 µm. (D-F) Microtubule organization. (D) ApPr from GFP^+^ CSF-cNs (green) protrude to the central canal and show fibrillary processes (arrowheads). (E) Acetylated tubulin (AcT, yellow) staining shows a dense network of stable microtubules at the ApPr and tightly packed cilia in the central canal. (F) γ-Tubulin staining (γTub, blue) reveals basal bodies (arrows) associated with ependymal cilia, but not with the GFP^+^ ApPr or their fibrillary protrusions (arrowheads). See also Videos S1 and S2. Scale bar, 3 µm. (G-I) Genetic absence of motile cilia on CSF-cNs. (G) In *GATA3^eGFP^*; *FoxJ1*-*CreER*-*tdTomato* mice, CSF-cNs stain green. (H) CSF-cNs are tdTomato-negative (tdT, magenta), confirming the absence of motile cilia. (I) GFP^+^ cells also express the CSF-cN receptor PKD2L1 at their core and protrusions (cyan). Scale bar, 20 µm. (J-L) Close apposition of ependymal cilia to the ApPr. (J) Overview showing GFP^+^ ApPrs protruding into the cc. (K) tdTomato^+^ ependymal cells (tdT, magenta) show their cilia (on transversal section, arrowheads) protruding into the cc and in close apposition to ApPrs. (L) AcT^+^ staining (yellow) shows cilia (arrows) from tdTomato^+^ ependymal cells, some of them in close proximity to ApPrs (arrowheads). Scale bar, 3 µm. (M) GFP^+^ ApPrs (green) are devoid of cilia, whereas tdTomato^+^ ependymocytes (tdT, magenta) display AcT^+^ motile cilia (AcT, yellow). Primary cilia are restricted to the soma. Adenylate cyclase III (AC3, cyan) labels primary cilia on the AcT-rich CSF-cN soma (inset, arrows). Scale bar, 10 µm; inset, 5 µm. (N, O) Ultrastructural confirmation. (N) Transmission electron microscopy (TEM) image showing a motile (9+2) cilium (green shading) and a putative primary (or the proximal segment of a motile, 9+0) cilium (magenta shading) adjacent to protrusions from the ApPr (yellow shading). Inset, a higher-magnification view reveals the ApPr membrane accommodating the surrounding cilia (Video S3). Scale bar, 1 µm; inset, 200 nm. (O) 3D reconstruction from 26 serial-section TEM images confirms cilia (green) are embedded in invaginations of the ApPr surface. The central canal lumen is outlined with a white dashed line (cc) in panels D and J.

We next analyzed these processes using an immunophenotypic approach. Staining for acetylated tubulin (AcT), a marker of stable microtubules in cilia,^19^ robustly labeled ependymal cilia but failed to label any fibrillary structures arising from GFP^+^ ApPrs (Figure 1D-F). Figure 1F shows a representative example of a long GFP^+^ process emanating from the ApPr (white arrowhead), completely devoid of AcT, which runs parallel to a AcT positive cilium. Similarly, staining for γ-tubulin (Figure 1F, S1C) and pericentrin (Figure S4; Videos S1 and S2) showed basal bodies adjacent to the cc that were not associated with the ApPr of CSF-cNs, where centrosomes were expected to be located. To genetically confirm the absence of motile cilia,^20,21^ we crossed *GATA3^eGFP^*with *FoxJ1*-*CreER*-*tdTomato* mice to differentially label ependymal cells and CSF-cNs. FoxJ1 is a transcription factor involved in ciliogenesis,^22,23^ however, we found no co-expression of tdTomato in GFP^+^ CSF-cNs, confirming these neurons are not ciliated (Figure 1G-I). Interestingly, tdTomato^+^ cilia from ependymal cells established close contacts with the ApPrs (Figure 1J-L, arrows). Staining for adenylate cyclase 3 (AC3), a marker of primary cilia, confirmed that although CSF-cN cell bodies possess an internalized primary cilium,^9^ their ApPrs are devoid of cilia (Figure 1M). Finally, we turned to transmission electron microscopy (TEM) with 3D reconstruction. Figure 1N is a high magnification EM image of an ApPr (highlighted in yellow) and Figure 1O the corresponding serial 3D reconstruction. The ApPr membrane was often deformed to accommodate the cilia of adjacent ependymal cells (Figure 1N, inset; Video S3), explaining the close spatial relationship observed by light microscopy (Figure S1C). This ultrastructural analysis unequivocally demonstrated that mouse CSF-cN ApPrs lack cilia.

Our results concur with previous SBF-SEM and TEM data which did not find cilia on the ApPr of CSF-cN from mice, but instead some protrusions that were referred to by the authors as cilia-like structures.^24^

### ApPrs are anatomically and biochemically segregated compartments

Having established that mouse ApPrs lack cilia, we next asked what structural specializations define this unique neuronal compartment.

The ApPr protrudes into the cc through a constricted neck that is surrounded by actomyosin complexes of adjacent ependymal cells (Figure 2A-D). This interphase is stabilized by adherens and tight junctions, as revealed by scanning (Figure 2 E-F) and transmission (Figure 2G) electron microscopy and ZO-1 immunostaining (Figure 2H). 3D reconstruction further showed an accumulation of cytosolic GFP at this neck (Figure S2), consistent with a role as a diffusion barrier to maintain the ApPr within a distinct biochemical environment. Indeed, the ApPr is located in a different extracellular environment than the rest of the cell, and these anatomical specializations probably play a very important role in the functional ^17^ and anatomical segregation of the ApPr.

**Figure 2.**
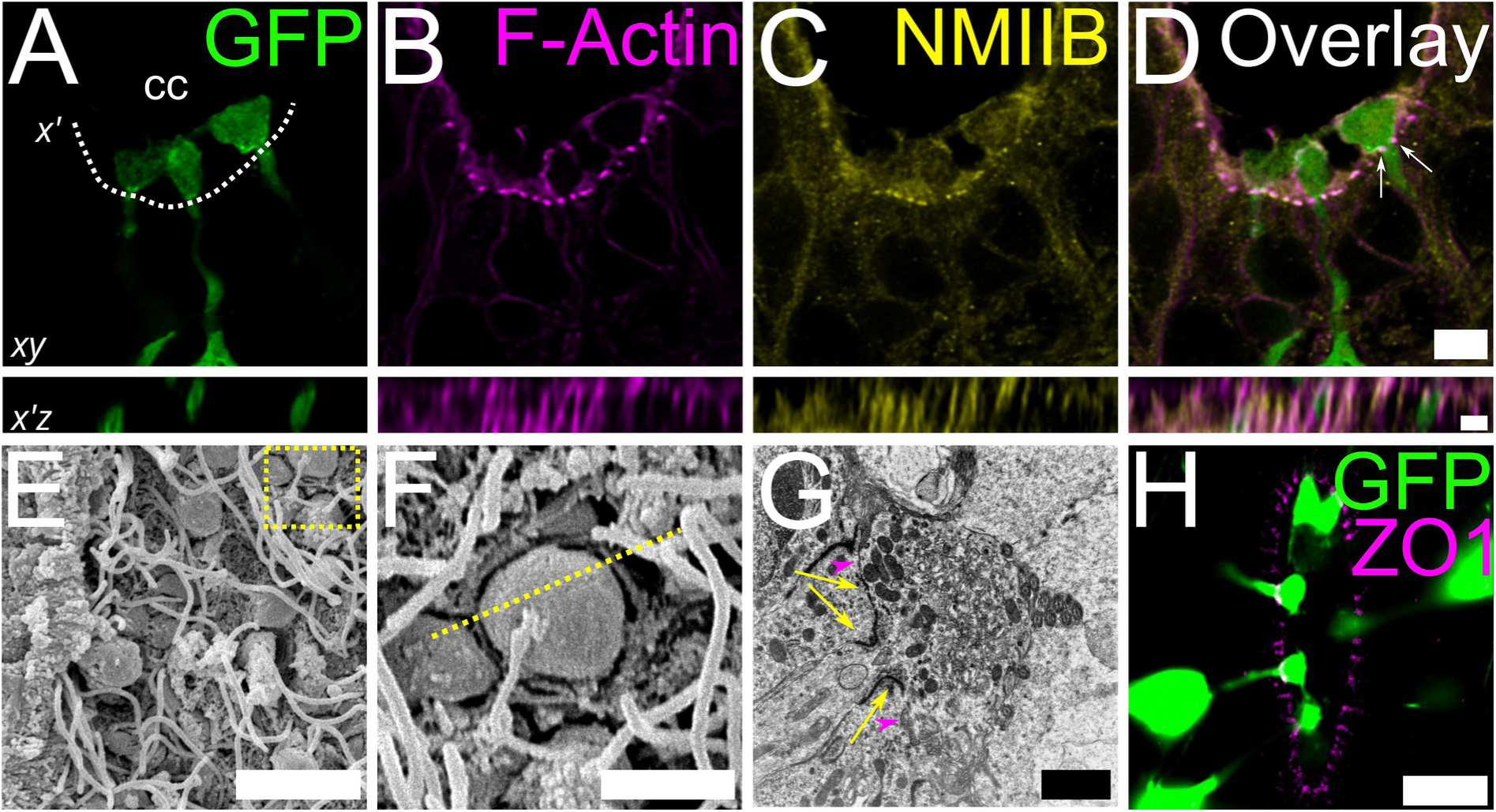
The ApPr is a structurally segregated compartment at the central canal. (A) Localization of the ApPr. A single confocal plane (top) and an x’z reslice (bottom, from 33 z-slices) show CSF-cN apical processes (ApPrs, green) projecting into the cc. The dashed line indicates the x’ plane, at the ependymo-luminal interphase. (B-D) Cytoskeletal specialization at the ApPr interphase. (B) Phalloidin staining reveals F-actin enrichment in ependymal cells and at the ApPr cortex. (C) Non-muscle myosin IIB is similarly enriched at these sites. (D) Overlay of (B) and (C), highlighting the actomyosin complexes surrounding the ApPr neck (arrows). Scale bar, 5 µm (xy), 2 µm (x’z). (E, F) Scanning electron microscopy (SEM) of the cc in horizontal section. (E) A panoramic view shows the ependymal wall (left) and multiple ApPrs protruding among a forest of microvilli and cilia. The yellow dotted square indicates the region magnified in (F). Scale bar, 5 µm. (F) A top-down view of an ApPr reveals its distinct, bulbous morphology amidst numerous, uniform ependymal cilia. The yellow dotted line indicates the approximate plane of section for (G). Scale bar, 2 µm. (G) Ultrastructural evidence of junctional complexes. Transmission electron microscopy (TEM) of an ApPr stem shows electron-dense adherens junctions (yellow arrows) and tight junctions (magenta arrowheads) anchoring it to adjacent ependymal cells. Scale bar, 500 nm. (H) Confirmation of tight junctions. Immunostaining for ZO-1 (magenta) at the base of GFP^+^ ApPrs (green) confirms the presence of tight junctions. Scale bar, 10 µm. The central canal lumen is outlined with a white dashed line (cc) in panel A.

### A drebrin- and doublecortin-stabilized cytoskeleton defines the ApPr and its protrusions

We next aimed to characterize the ApPr cytoskeleton. Immunohistochemistry for filamentous F-actin, a highly dynamic cytoskeletal protein, revealed that the ApPr protrusions extending into the canal lumen are also highly enriched in F-actin (Figure 3A-C).

**Figure 3.**
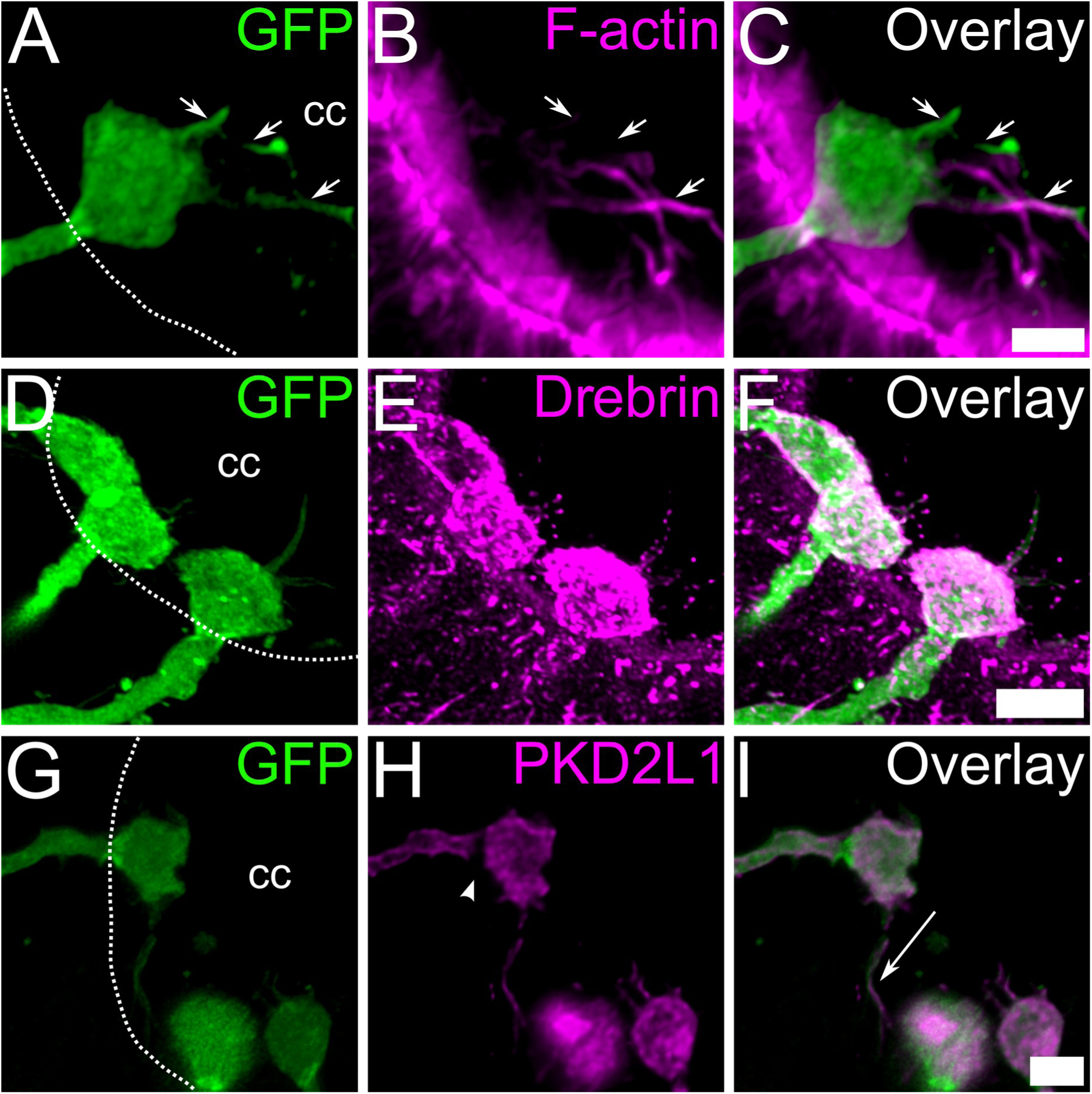
The CSF-cN apical process is a drebrin-stabilized compartment that emits lumen-protruding structures. (A) A GFP^+^ CSF-cN apical process (ApPr) extends into the cc and emits filamentous protrusions (arrows). (B, C) The ApPr and its protrusions are actin-rich. (B) Phalloidin staining (magenta) reveals F-actin in ependymal microvilli and in long, filamentous structures within the lumen. (C) Overlay confirms these F-actin-rich structures (arrows) originate from the GFP^+^ ApPr. Maximum intensity projection of 39 confocal planes. Scale bar, 3 µm. (D) High-magnification view showing the complex, tortuous surface of the ApPr, with instances of contact between neighboring ApPrs. (E, F) Drebrin enrichment stabilizes the ApPr. (E) Immunostaining shows strong drebrin (magenta) localization throughout the ApPr. (F) Overlay reveals drebrin-positive puncta along the protrusions (see also Video S4). Maximum intensity projection of 55 confocal planes. Scale bar, 5 µm. (G) A diversity of protrusions emanate from ApPrs, including some that appear to connect distant ApPrs (see also Video S5). (H) Compartmentalized PKD2L1 expression. PKD2L1 (magenta) is highly enriched on the ApPr surface but is reduced at the constricted neck region (arrowhead). (I) ApPr protrusions interact with the ependymal surface. A GFP^+^/PKD2L1^+^ protrusion (arrow) extends along the apical surface of a tdTomato^+^ ependymal cell (gray) in a *GATA3^eGFP^*; *FoxJ1*-*CreER*-*tdTomato* mouse. Maximum intensity projection of 19 confocal planes. Scale bar, 3 µm. The central canal lumen is outlined with a white dashed line (cc) in panels A, D and G.

The ApPr core also exhibits a stable microtubule network, as revealed by conspicuous AcT staining (Figures 1E, 1L). Then we analyzed the microtubule-associated protein doublecortin (DCX), an immature neuron marker known be expressed in CSF-cNs,^25^ and—more generally—to regulate microtubule structure in neuron morphology and plasticity.^26,27^ DCX immunostaining also revealed stable microtubule networks in ApPr cores (Figure S1A-C), consistent with their acetylated tubulin enrichment (Figure S3), an organization reminiscent of dynamic structures like growth cones.^28,29^ DCX is expressed in migrating neurons,^28^ and DCX-bound microtubule bundles converge at the centrosome,^30^ although in CSF-cNs ApPrs DCX did not colocalize with basal bodies (Figures S1C, S6 and Videos S1, S2). The co-localization of DCX with the stable microtubule cytoskeleton within the ApPr supports a role in reinforcing this dendritic specialization

Since the ApPr is a dendritic specialization enriched in F-actin and stable tubulin that also shares ultrastructural features with dendritic spines,^31^ we investigated the presence of drebrin 1 (DBN1). Drebrin is an actin-, tubulin-binding protein that accumulates in dendritic spines, where it plays a role in spine remodeling during potentiation and synapse formation.^32–36^ We found that the ApPr itself is stabilized by a drebrin-rich, specialized cytoskeleton (Figure 3D-F). Notably, drebrin is also present in the ApPr protrusions (Video S4) but is much less abundant in the soma (not shown). These protrusions also stain positive for the multimodal sensor PKD2L1 (Figure 3G-H). Unlike GFP, PKD2L1 staining is weaker at the ApPr neck, indicating some form of spatial segregation (Figure 3H). Finally, these protrusions project either centripetally into the canal or run over the surface of td-Tomato^+^ ependymocytes and seem to contact more distant ApPrs (Figures 3I, S4; Video S5).

### The ApPr is a complex compartment with filopodia and specialized membrane domains

Given their lack of cilia, we therefore studied the ApPr protrusions in detail. These numerous protrusions radiate (Figure 4A, arrows) from the drebrin-rich ApPr core (Figure 4B-C), resembling filopodia and extending in all directions with varying lengths and widths (Figures 4D, S4A). Supravital staining with the nanoenvironmental probe LAURDAN^37^ reveals that the ApPr core (Figure 4E) is lipid-rich (Figure 1F), and phasor-based hyperspectral analysis ^38^ shows that both the ApPr core and its protrusions are enriched in gel-phase lipids (Figure 4E, 4G-G’), indicating that their plasma membrane exhibit lipid rafts or less fluid regions from which protrusions seem to arise. As drebrin has been shown to mediate actin clustering in neuronal filopodia,^36^ we analyzed the presence of the filopodial marker myosin-X.^39–42^ We found that the major actin-rich protrusions arising from the ApPr core stain positive for myosin-X, indicating these are indeed filopodia (Figure 4I-L; Video S6).

**Figure 4.**
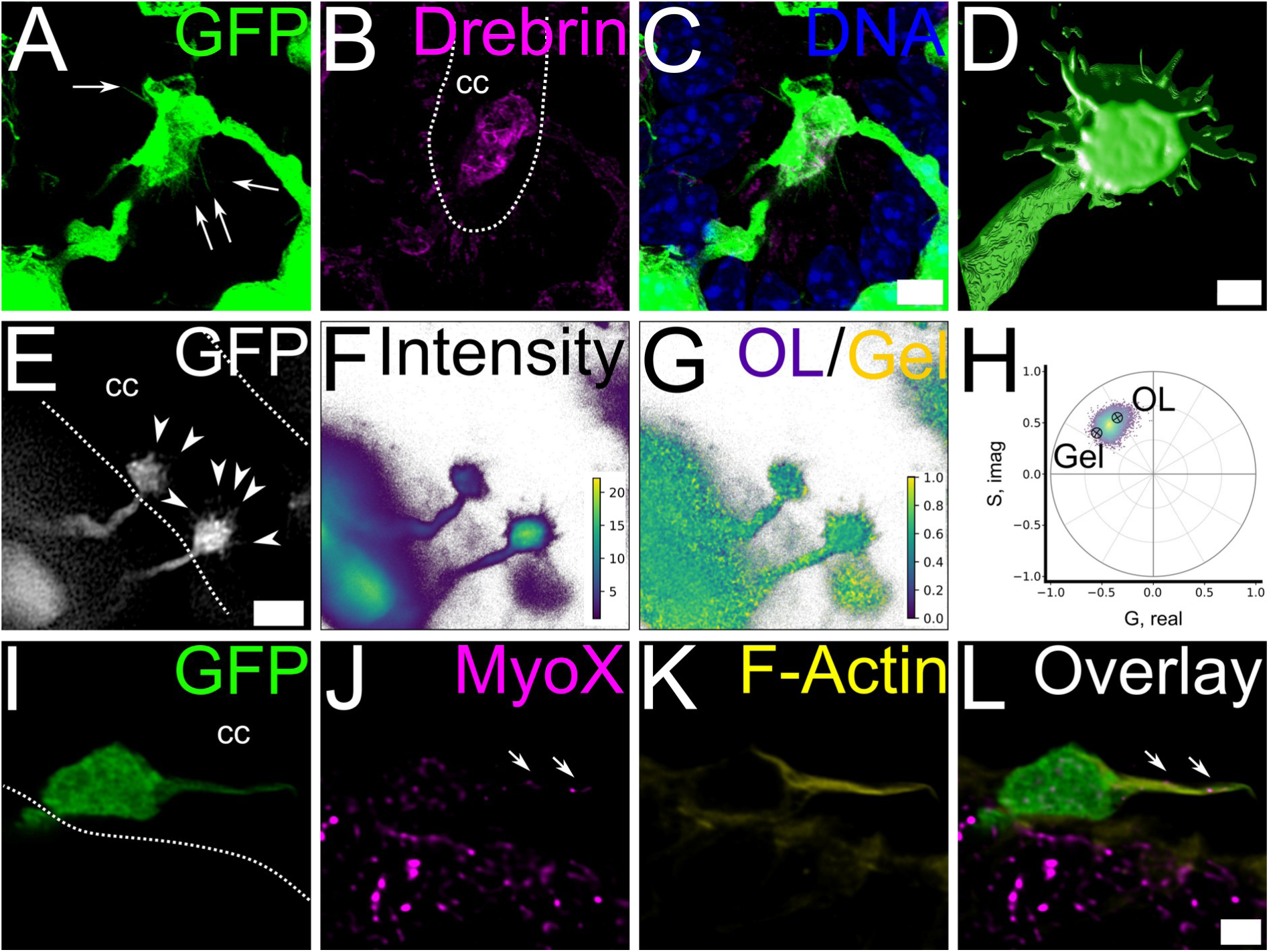
The ApPr plasma membrane is organized into rigid, gel-phase lipid microdomains from which filopodia arise. (A-C) Protrusions originate from a drebrin-stabilized core. (A) Multiple filiform projections (arrowheads) emerge from a cluster of GFP^+^ ApPrs. (B) Immunostaining reveals strong drebrin enrichment within the ApPrs. (C) Overlay shows the projections emanating from the drebrin-rich core (maximum intensity projection of 48 confocal planes). Scale bar, 5 µm. (D) Surface reconstruction of a GFP^+^ ApPr illustrates its bulbous shape and numerous surface protrusions. Scale bar, 2 µm. (E) Live imaging in a *GATA3^eGFP^* (gray) spinal cord slice (oblique section) confirms the abundance of these radiating projections (arrows). Scale bar, 5 µm. (F, G) The ApPr membrane is enriched in gel-phase lipid microdomains. (F) LAURDAN intensity in live tissue shows high incorporation in the ApPr core. Color gradient shows the intensity from low (blue) to high (yellow). (G) Pseudo-colored phasor analysis of LAURDAN hyperspectral imaging maps membrane fluidity, revealing rigid, gel-phase lipid microdomains (warmer colors) that are prominent across the ApPr surface and its protrusions. Color gradient shows lipid packing from low (blue) to high (yellow). (H) The corresponding phasor plot, where the red cursor indicates the gel-phase component fraction represented in (G). Color gradient shows pixel density from low (blue) to high (yellow). (I-L) Protrusions arising from the ApPr core are filopodia. (I) The ApPr emits major GFP^+^ protrusions into the cc (green). (J) Immunostaining reveals both the ApPr and its protrusions (arrows) are positive for the filopodial marker Myosin-X (magenta). (K) Phalloidin staining shows these structures are enriched in F-Actin (yellow). (L) Overlay shows the actin-enriched protrusion is a filopodia (arrows; Video S6). Scale bar, 2 µm.The central canal lumen is outlined with a white dashed line (cc) in panels B, E and I.

Ultrastructural analysis of the ApPr using transmission electron microscopy revealed that the ApPr is a complex structure containing mitochondria, vesicles, multivesicular bodies, and an extensive system of tubules and cisternae (Figures 1N, 2G). ApPrs stain positive for the ER marker calnexin (Figure 5A-B), and for lysosome-associated membrane protein-1 (LAMP1, Figure 5C-D), indicating active membrane dynamics and a prominent endocytic apparatus within the ApPr. Immunostaining reveals that some ApPrs exhibit PKD2L1^+^ protrusions (Figure 3G-I), as previous evidence suggested,^17,43^ while others show a central accumulation of the receptor (Figure 5E-G), which may indicate local biosynthesis, recycling or a robust anchorage system. Transmission electron microscopy also revealed omega figures at the plasma membrane (Figure 5H, arrowheads), consistent with endocytic processes. Additionally, caveolin-3 (Cav3) immunostaining confirmed the presence of caveolae at the ApPr (Figure 5I-L), indicating that ApPrs are capable of clathrin-independent endocytosis.

**Figure 5.**
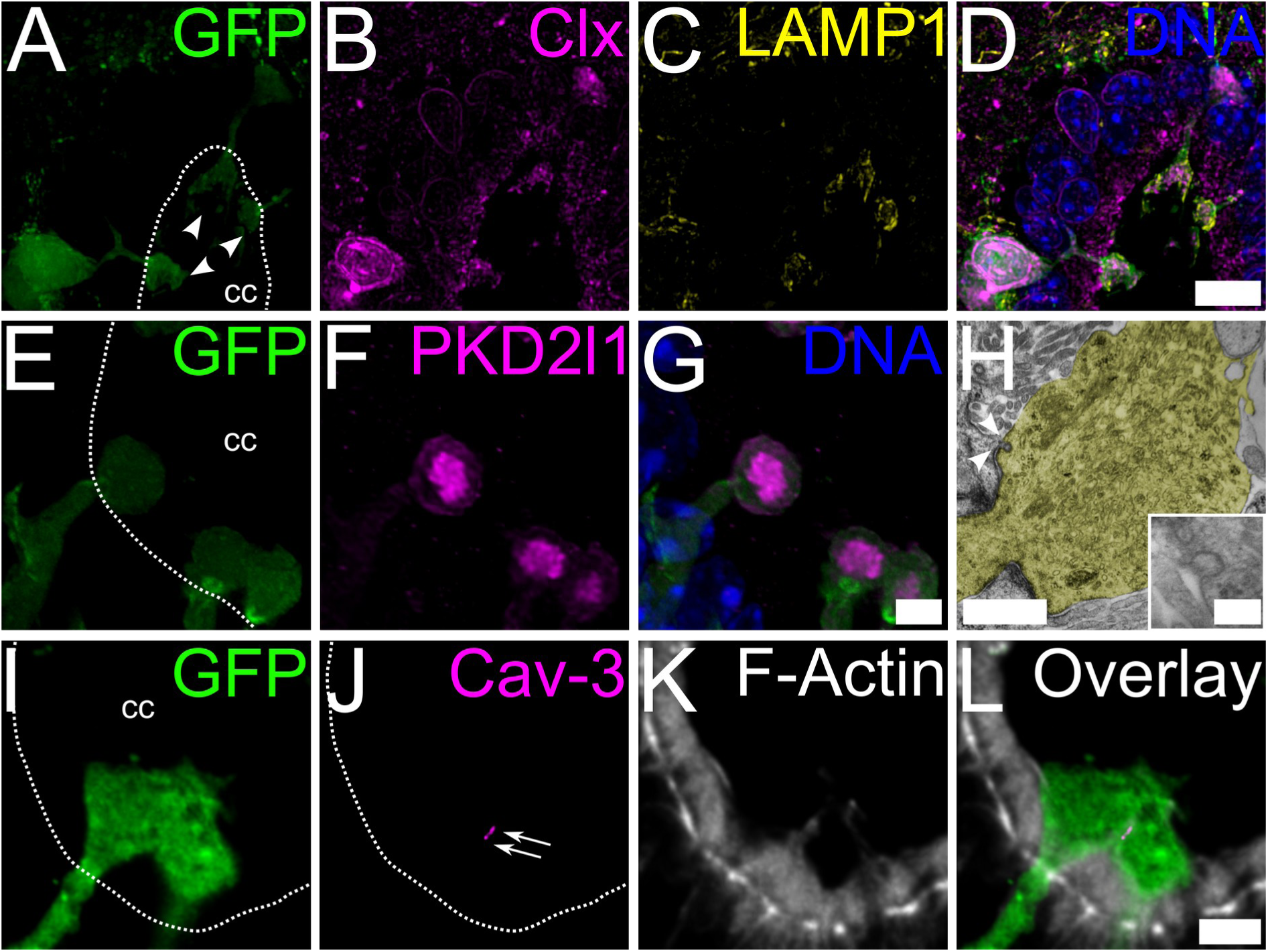
The ApPr is an autonomous compartment with a specialized endomembrane system. The ApPr contains organelles for membrane trafficking and recycling or degradation. (A) Three GFP^+^ ApPrs projecting into the cc (arrowheads). (B) The same ApPrs immunostained for the endoplasmic reticulum marker calnexin (CLX, magenta). (C) The ApPr periphery is enriched with the lysosomal marker LAMP1 (yellow). (D) Overlay of A, B, and C (maximum intensity projection of 30 confocal planes). Scale bar, 10 µm. (E-G) PKD2L1 channels accumulate in the ApPr core. Intense PKD2L1 immunostaining (magenta) is observed in the central region of some ApPrs (green), suggesting local clustering or biosynthesis. Scale bar, 3 µm. (H) Ultrastructural evidence of active membrane dynamics. Transmission electron microscopy of an ApPr (yellow shading) reveals membrane invaginations (white arrowheads) indicative of active endocytosis. Inset, a higher magnification view shows a caveola-like structure. Scale bar, 1 µm; inset, 100 nm. (I-L) Caveolae are present at the ApPr plasma membrane. (I) Juxtaposed GFP^+^ ApPrs (green). (J) Immunostaining for caveolin-3 (Cav3, magenta) reveals punctate staining (arrows). (K) Phalloidin staining (gray). (L) Overlay reveals F-actin at the proximity of Cav3 localization on the contacting ApPr surfaces. Scale bar, 3 µm. The central canal lumen is outlined with a white dashed line (cc) in panels A, E, I and J.

### Direct mechanostimulation of the cilia-free ApPr evokes PKD2L1-dependent currents and action potentials

We next investigated if the ApPr could still be a mechanosensory compartment with a mechanism divergent from the ciliary model.^5^ Therefore, we tested the mechanosensory capability of the cilia-free mouse ApPr.

To probe the intrinsic mechanosensitivity of the ApPr, we performed patch clamp recordings from CSF-cN in acute, adult spinal cord slices at near physiological temperature, and used a second blunt pipette for mechanical stimulation. A 2 µm indentation (push) of the ApPr reliably evoked phasic inward currents that subsided rapidly (−43.7 ± 23.6 pA, n = 15 cells), suggesting some form of desensitization (Figure 6A).

**Figure 6.**
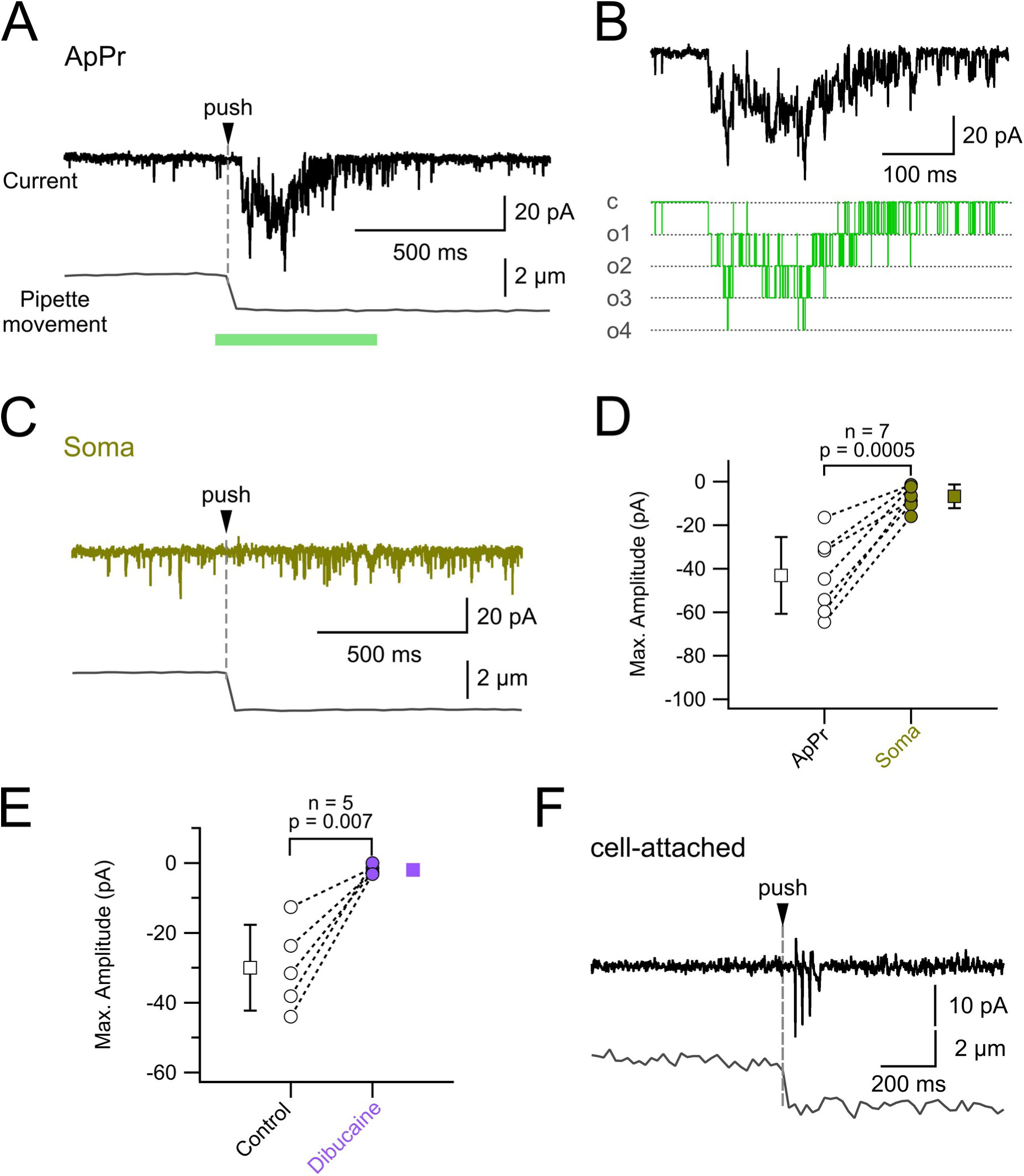
The cilia-free ApPr is a direct, PKD2L1-dependent mechanosensor. (A) Representative whole-cell voltage-clamp recording from a CSF-cN ApPr showing inward currents evoked by mechanical stimulation of the ApPr (2 µm indentation, “push”; arrowhead). The lower black trace show stimulus pipette position. The horizontal green line indicates the region magnified in B. Scale bars, vertical, 20 pA, 2 µm; horizontal, 500 ms. (B) Example of mechanically-evoked currents (upper, black trace) and the corresponding idealized events (lower, green trace) indicating the different states of the channels: closed (c), one (o1), two (o2), three (o3) or four (o4) open channels. The traces correspond to the region indicated with the green horizontal line in A. Scale bars, vertical, 20 pA; horizontal, 100 ms. (C) Representative recording from the same neuron as in A showing that identical mechanical stimulation of the soma evokes negligible current responses (olive trace). The lower black trace shows the stimulus pipette position. (D) Quantification of the maximum current amplitude evoked by mechanical stimulation at the ApPr (black) versus the soma (olive). Each circle represents one cell; squares and bars indicate mean ± SD. ApPr responses are significantly larger (p = 0.0005, two-sided paired t-test, n = 7 cells, effect size = 2.79, statistical power = 0.999). (E) The PKD2L1 channel inhibitor dibucaine abolishes mechanosensitive currents at the ApPr. Quantification of the maximum current amplitude evoked by mechanical stimulation before (Control, black) and after dibucaine application (Dibucaine, violet) shows a significant reduction (p = 0.007, one-sided paired t-test, n = 5 cells, effect size = 3.20, statistical power = 0.999). In panels B-D, circles represent individual cells; filled squares and error bars represent mean ± SD. (F) Mechanical stimulation of the ApPr evokes firing in a representative cell-attached recording (black trace). The lower black trace shows stimulus pipette position.

A close inspection of the current responses induced by mechanical stimulation clearly shows single-channel activity that is characteristic of PKD2L1-mediated currents in CSF-cNs (Figure 6B).^7,16,44^

We next tested whether a similar current could be recorded when mechanically stimulating the soma. As expected from previous data that indicate that PKD2L1 channels are exclusively located at the ApPr, identical or even stronger (4 µm pushes) mechanical stimulation of the soma evoked only minimal responses (−6.71 ± 5.45 pA; n = 7 cells, 5 animals) (Figure 6C) that were significantly smaller than ApPr-evoked ones (−43.10 ± 17.63 pA; p = 0.0005; paired t-test, n = 7 cells, 5 animals) and indistinguishable from baseline (Figure 6C, D).

In order to directly assess the involvement of PKD2L1 channels we performed the same experiments in the continuous presence of the PKD2L1 inhibitor dibucaine (200 µM). In these conditions, the ApPr mechanically-induced currents were almost abolished (mean amplitude: −30.0 ± 12.3 pA vs. −2.02 ± 1.27 pA; p = 0.007, paired t-test, n = 5 cells, 2 animals) (Figure 6E).^23,29^

To assess the functional output of this transduction, we performed cell-attached recordings. Repeated mechanical stimuli to the ApPr reliably evoked action potentials, demonstrating that mechanosensitive currents are sufficient to drive neuronal firing (n = 4 ells, 3 animals)(Figure 6F).

Taken together, these results establish that direct mechanical stimulation of the cilia-free ApPr activates resident PKD2L1 channels to generate depolarizing currents sufficient to trigger action potentials.

## DISCUSSION

Our study challenges the prevailing model of mechanosensation in cerebrospinal fluid-contacting neurons (CSF-cNs) by identifying a fundamental evolutionary divergence. We demonstrate that, in contrast to their ciliated counterparts in zebrafish,^6,7^ mouse CSF-cN apical processes (ApPrs) are devoid of cilia. This conclusion, supported by ultrastructural analysis, transgenic labeling, and immunophenotyping, suggest the need of a re-evaluation about how these spinal sensory cells detect mechanical stimuli. The preservation of PKD2L1 expression^45^ and mechanosensory function in the absence of a ciliary apparatus points to a novel, cilia-independent transduction mechanism in mammals.

The close apposition of ependymal cilia to the ApPr surface could explain confounding data.^15,18,25^ However, our high-resolution data unequivocally demonstrate that the irregular protrusions emanating from the mouse ApPr are not cilia. Primary cilia are present only in the soma, as previously described^9^ and might underlie the impaired motor function observed in PKD2L1-driven cilia ablation experiments.^15^

Neurons, like other differentiated cells, feature distinct, actin-rich, finger-like extensions known as filopodia and filopodia-like protrusions. These protrusions play a role in environmental sensing and cell migration.^46^ Arising from the dense cortical actin meshwork, they are highly dynamic structures that continuously remodel in response to environmental cues.^47^ Although widely studied in developmental processes, the functional role of dendritic filopodia during adulthood has only recently begun to be uncovered.^48,49^

We identified a specialized cytoskeletal and membrane organization that underlies this cilia-independent mechanosensation. The ApPr core is enriched with drebrin-stabilized actin filaments and a doublecortin-positive microtubule network, creating a rigid,^28,29^ compartmentalized structure whose neck likely acts as a diffusion barrier. Furthermore, we show that the ApPr membrane is organized into gel-phase lipid microdomains—a biophysical state well-suited for mechanotransduction, as the lipid bilayer mediates mechanical force transmission and directly influences mechanosensitive channel gating.^50,51^ This unique architecture may anchor PKD2L1 channels and enables efficient transmission of mechanical force, eliminating the need for the ciliary structure.

From these microdomains, actin-rich, drebrin-stabilized PKD2L1-positive filopodia, filopodia-like structures^24,52^ and lamellae^53,54^ arise. These might represent the genuine receptorial sensory specializations. Phosphoinositide lipids can pack densely into highly dynamic microdomains regulating membrane organization and exo- and endocytosis,^55–57^ and have been shown to nucleate and force proteins to protrude into filopodia-like structures.^47,58^

We have recently shown that pH-sensitivity in CSF-cNs is similarly compartmentalized to the ApPr,^17^ suggesting a conserved principle: spatial and functional segregation of sensory modalities to a specialized dendritic subdomain. Their extension into the cc lumen positions them as ideal candidates for directly sensing CSF dynamics. In addition, the intimate spatial relationship with the motile cilia of ependymal cell brings about the possibility that CSFcNs may sense the motion of the CSF and/or ciliary beat.

Beyond detecting transient stimuli, the ApPr sensory apparatus allows for continuous monitoring of the CSF environment. The PKD2L1-dependent persistent current we described^17^ suggests that CSF-cNs can encode both phasic mechanical events and sustained shifts in CSF composition. Indeed, this compartment has proven to be highly sensitive to pH shifts near physiological values.^17^ This supports the view that CSF-cNs are multimodal integrators that convey chemical and mechanical information to spinal motor circuits, potentially modulating locomotion and posture,^5,7^ and even contributing to signaling processes involved in spinal cord repair.^59^

In conclusion, we have defined a cilia-independent paradigm for mechanosensation in spinal sensory neurons in mice. The mouse CSF-cN ApPr is a sensory compartment that employs a drebrin-stabilized cytoskeleton and specialized membrane domains to directly transduce mechanical forces via PKD2L1 channels. This evolutionary innovation underscores the plasticity of cellular sensory mechanisms and redefines our understanding of how the spinal cord monitors its internal environment. The molecular machinery we describe opens a new path for exploring the role of CSF-cNs in sensorimotor integration and even related to neurological disorders.

### Limitations of the study

Several important questions remain. Our *ex vivo* assay demonstrates the intrinsic mechanosensitivity of the ApPr, though the nature of the relevant physiological stimulus *in vivo*—be it CSF flow, pressure, or spinal movement—is still unknown. Finally, the precise molecular mechanisms linking the specialized cytoskeleton and membrane domains to PKD2L1 gating await definition. Addressing these points will provide the key to fully understanding the role of this sensory compartment.

## METHOD DETAILS

### Experimental model and subject details

Juvenile mice (P20-P30) of both sexes from the following established lines were used: *GATA-3^eGFP^*,^60^ *FOXJ1-CreER*-R26R*tdTomato*.^61^ Double trangenics were obtained by standard mating procedures. We induced the expression of tdTomato by intraperitoneal injection of tamoxifen (Sigma Millipore; 2 mg, 20 mg/ml in corn oil) for 5 consecutive days and allowed a 10-day clearance period before experiments. All animal procedures were approved by our local Committee for Animal Care (CEUA 001/01/2022a). Mice were housed at 21–22℃ and 50%-60% humidity on a 12/12 h light cycle in cages (30 × 20 cm) containing < 6 animals. Cages were cleaned twice weekly and enriched with cardboard cylinders.

### Slice preparation and electrophysiology

We prepared transverse or 45°-angled lumbar spinal cord slices (300 µm thick) from GATA-3^eGFP^ mice as previously described.^17^ Briefly, under isoflurane anesthesia, we decapitated mice and rapidly dissected the spinal cord in ice-cold, oxygenated dissection Ringer (in mM: 101 NaCl, 3.8 KCl, 1.3 MgSO₄·7H₂O, 1.2 KH₂PO₄, 10 HEPES, 25 glucose, 1 CaCl₂, 18.7 MgCl₂, pH 7.4, 300 mOsm/kg H₂O). After laminectomy, we embedded the cord in 4% low-melting-point agarose for slicing.

We performed patch-clamp recordings from CSF-cNs in submerged slices maintained at 34 ± 1°C (Peltier system; Luigs & Neumann) under an upright Olympus microscope (60×, 1.0 NA objective). The extracellular recording solution contained (in mM): 115 NaCl, 2.5 KCl, 1.3 NaH₂PO₄, 26 NaHCO₃, 25 glucose, 5 Na-pyruvate, 2 CaCl₂, 1 MgCl₂, pH 7.4 (continuously bubbled with 95% O₂/5% CO₂, 300 mOsm/kg H₂O). The intracellular solution contained (in mM): 165 K-gluconate, 10 HEPES, 1 EGTA, 0.1 CaCl₂, 4.6 MgCl₂, 4 Na₂-ATP, 0.4 Na-GTP, 0.04 Alexa 594, pH 7.3, 300 mOsm/kg H₂O. Pipette resistance was 6–7 MΩ; series resistance was monitored but not compensated.

We used a dual-pipette approach: one pipette for whole-cell or cell-attached recording and a second, independent pipette for mechanical stimulation. We positioned pipettes with Luigs & Neumann micromanipulators. To stimulate the ApPr or soma, we used the micromanipulator’s ‘step’ function to deliver precise 2 µm (or 4 µm) indentations (push).

We acquired data at 20 kHz (low-pass filtered at 2.9 kHz) using a Heka Elektronik EPC 10 USB Patch Clamp Amplifier (RRID:SCR_018399) and Patchmaster software (RRID:SCR_000034). We visualized eGFP and Alexa 594 using a dual LED light source (Optoled; Cairn Research) and an Andor Ixon EM CCD camera (Oxford Instruments) with appropriate filter sets (Chroma, USA).

### Pharmacology

We applied the PKD2L1 channel inhibitor dibucaine (Sigma-Aldrich Cat# D0638) by puffing a 200 µM solution in extracellular Ringer for 30 s.

### Electrophysiological analysis

We analyzed electrophysiological data in IGOR Pro (WaveMetrics, USA; RRID:SCR_000325) using NeuroMatic (RRID:SCR_004186),^62^ TaroTools^63^ and custom routines.

### Morphological identification of patch-clamp recorded cells

During whole-cell recordings, we filled cells with biocytin via the patch pipette. After recording (15–30 min), we fixed slices by immersion in 4% formaldehyde from PFA in 0.1 M PB, pH 7.4, for 1 h and processed them for immunohistochemistry as previously described.^64^

### Immunohistochemistry

For immunohistochemistry, we anesthetized mice by administering intraperitoneally (in mg/kg): 100 ketamine, 10 xylazine, 10 diazepam, and fixed them by intracardiac perfusion with 4% formaldehyde from PFA in 0.1 M PB. We sectioned spinal cords (50–70 µm) on a vibrating microtome, blocked sections in PBS with 0.5% BSA (30 min), and incubated them overnight with primary antibodies in PBS containing 0.3% Triton X-100. After washing, we incubated sections with fluorophore-conjugated secondary antibodies and mounted them in glycerol-based medium.

The following primary antibodies were used: mouse anti-drebrin (Enzo Life Sciences Cat# ADI-NBA-110, RRID:AB_10621721; 1:100), rabbit anti-calnexin (Abcam Cat# ab22595, RRID:AB_2069006; 1:1000), goat anti-doublecortin (Santa Cruz Biotechnology Cat# sc-271390, RRID:AB_10610966; 1:100), mouse anti-acetylated tubulin (Sigma-Aldrich Cat# T7451, RRID:AB_609894; 1:1000), rabbit anti-gamma-tubulin (Sigma-Aldrich Cat# T5192, RRID:AB_261690; 1:1000), rabbit anti-adenylate cyclase type 3 (Santa Cruz Biotechnology Cat# sc-588, RRID:AB_630839; 1:50), rabbit anti-myosin-X (Invitrogen, PA5-76391, 1:50-1:1000), rabbit anti-PKD2L1 (Millipore Cat# AB9084, RRID:AB_571091; 1:500), mouse anti-non muscular myosin 2b (DSHB Cat# CMII 23, RRID:AB_528359; 1:20), rabbit anti-pericentrin (5 µg/ml),^65^ rabbit anti-ZO1 (Thermo Fisher Scientific Cat# 61-7300, RRID:AB_2533938; 1 µg/ml), mouse anti-LAMP1 (DSHB Cat# 1d4b, RRID:AB_2134500; 1:10), mouse anti-Caveolin 3 (Santa Cruz Biotechnology Cat# sc-5310, RRID:AB_626814; 1:50), F-Actin was stained with Alexa 647 conjugated phalloidin (Thermo-Fisher) according to the manufacturer’s specifications, DNA was stained either with 1µg/ml Hoechst 33342 (Thermos Fisher Cat# H1399) or 2 g/ml Methyl Green (Sigma-Aldrich Cat# 323829).^66^

### Confocal microscopy and image processing

We acquired images on a Zeiss LSM 800 with Airy scan confocal microscope (RRID:SCR_015963) using a 63× oil immersion lens (NA 1.4). We set sampling intervals according to the Nyquist criterion. We deconvolved z-stacks using Huygens Essential 4.5 (RRID:SCR_014237) with an experimental PSF and the Classic Maximum Likelihood Estimation algorithm (40 iterations, quality threshold 0.05). We assembled composites and adjusted brightness/contrast minimally in FIJI/ImageJ (RRID:SCR_002285). We assembled figures using Inkscape 1.4.2 (RRID:SCR_014479).

### Hyperspectral imaging (HSI) of lipid fluidity

To assess membrane fluidity, we incubated live spinal cord slices (300 µm) with 3 µM LAURDAN (1-[6-(dimethylamino)naphthalen-2-yl]dodecan-1-one) for 15 min in the extracellular Ringer. We acquired fluorescence spectra in xyλ-mode using the Zeiss LSM 800. We performed spectral phasor analysis using PhasorPy v.0.7;^67^ v.0.750; the custom script is available at http://codeberg.org/dprieto/CSF-cN2026.

### Electron microscopy

For transmission electron microscopy (TEM), we fixed the spinal cord by perfusion with a mixture containing 4% formaldehyde (from PFA) and 1% glutaraldehyde in 0.1 M PB pH 7.4. The spinal cords were divided into 1 mm-thick segments, washed several times in PB, and post-fixed in 1% OsO₄, and epoxy resin embedding (Durcupan ACM, Sigma Aldrich). We located areas of interest on 1 μm-thick semi-thin sections stained with borax methylene blue, and then carved to obtain the final sectioning surfaces aiming to obtain series that would allow identification of structures in a three-dimensional context. We obtained ultrathin sections, contrasted with uranyl acetate and lead citrate, and imaged them with a JEOL JEM-100CX II TEM (RRID:SCR_020146) as previously described.^68^ We performed segmentation in TrakEM2^69^ and 3D reconstruction in FIJI.^70^

For scanning electron microscopy (SEM), we fixed horizontal and oblique vibratome spinal cord sections in 2.5% glutaraldehyde for 18 h, washed in 0.1 M PB pH 7.4 and dehydrated them in a graded ethanol series. We performed critical-point drying, sputter-coated samples with gold, and imaged them using a JEOL JSM-5900LV Scanning Electron Microscope as previously described.^71^

### Schematics and videos

We created schematics using Amadine 1.8 (BeLight Software, 2026) and Inkscape. We assembled supplementary videos using FIJI or Huygens software.

### Statistical analysis

We assessed normality using the Shapiro-Wilk test. We performed statistical analysis using IGOR Pro and R/RStudio. We defined *n* as the number of recorded cells. Data are presented as mean ± SD. For comparisons of paired data we used the paired t-test; we considered p<0.05 statistically significant. R source code is available at https://codeberg.org/dprieto/CSF-cN2026)

## RESOURCE AVAILABILITY

### Lead contact

Requests for further information and resources should be directed to and will be fulfilled by the lead contact, Daniel Prieto.

### Materials availability

Numerical data used to build the figures of this manuscript have been deposited at Redata (https://redata.anii.org.uy/). Artwork has been deposited at SciDraw (https://scidraw.io/).^72,73^

## ACKNOWLEDGMENTS

We thank Prof. Jonas Frisén for kindly sharing their FoxJ1-CreER-tdTomato mice; Drs. Paola Lepanto and José Badano for kindly sharing cilia-related antibodies and advice; Drs. Leonel Malacrida and Bruno Pannunzio for their valuable assistance with hyperspectral LAURDAN staining and PhasorPy coding; the late Prof. Stephen J. Doxsey for the anti-pericentrin antibody; Dr. Ana Laura Reyes-Ábalos for her assistance at the Scanning Electron Microscopy Unit, School of Sciences, Universidad de la República. The authors would also like to acknowledge B.S. Renata Simeone, and the staff of the Rodent Facility at Instituto de Investigaciones Biológicas Clemente Estable for assistance with mouse breeding and genotyping. The monoclonal antibodies CMII-23 developed by Conrad, G.W. & Conrad, A.H. and 1D4B developed by August, J.T. (Johns Hopkins School of Medicine) were obtained from the Developmental Studies Hybridoma Bank, created by the NICHD of the NIH and maintained at The University of Iowa, Department of Biology, Iowa City, IA 52242.

## AUTHOR CONTRIBUTIONS

Conceptualization, D.P., F.F.T., O.T.-C., R.E.R.; Methodology, D.P., F.F.T., O.T.-C., M.V.F.; Investigation, D.P., M.I.R., G.F., O.T.-C., M.V., F.F.T.; Formal analysis, F.F.T., M.V.; Data curation, F.F.T., D.P.; Visualization, D.P., M.C., F.F.T.; Writing – original draft, D.P.; Writing – review & editing, D.P., F.F.T., R.E.R.; Funding acquisition, R.E.R., F.F.T., D.P.; Resources, F.F.T., R.E.R.; Supervision, F.F.T., R.E.R.

## DECLARATION OF GENERATIVE AI AND AI-ASSISTED TECHNOLOGIES

During the preparation of this work the authors used Nature Research Assistant beta in order to copy-edit the manuscript. After using this tool or service, the authors reviewed and edited the content as needed and take full responsibility for the content of the publication.

## SUPPLEMENTAL INFORMATION INDEX

Legends for Videos S1-S5 in a PDF.

Video S1. Apical Processes of CSF-cNs are cilia-free and lack pericentrin-positive basal bodies. Three-dimensional projection. (Related to Figure 1 and S4).

Video S2. Confocal Z-Series showing exclusion of basal bodies from CSF-cN Apical Processes. (Related to Figure 1 and Video S1).

Video S3. Ultrastructural Evidence that the CSF-cN Apical Process is a cilia-free compartment. Transmission electron microscopy (TEM) serial-sections. (Related to Figure 1N).

Video S4. Drebrin enrichment in the Actin-cytoskeleton of CSF-cN Apical Processes. Deconvolved confocal Z-stack of the apical domain of CSF-cNs. (Related to Figure 3I and S4).

Video S5. 3D Reconstruction Reveals Interconnections Between Neighboring CSF-cN Apical Processes. (Related to Figures 3 and 4).

Video S6. Confocal Z-series showing Apical Process major protrusions are enriched in F-Actin and Myosin-X. (Related to Figure 4I-L).

**Figure S1.**
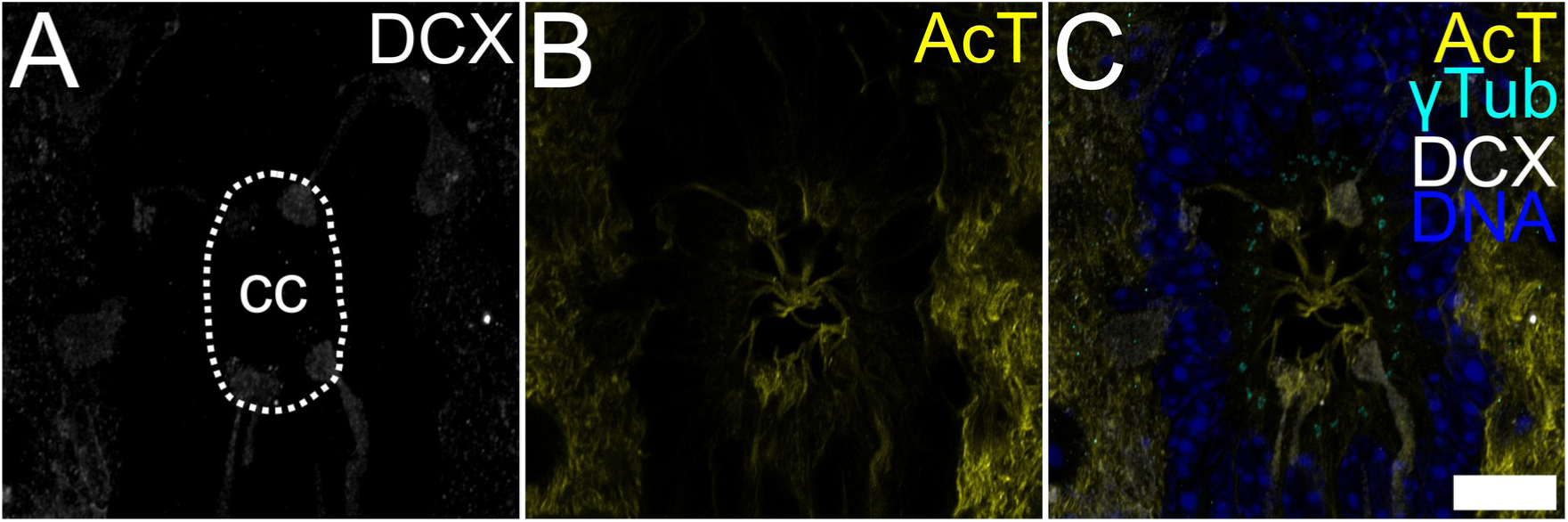
Mouse spinal cord central canal organization, related to Figure 1. (A) Topographical view of the central region of the mouse spinal cord in transverse section depicting the distribution of doublecortin-positive CSF-cN (DCX, gray) and their ApPrs protruding into the cc. (B) Stable microtubules including cilia stained with anti-acetylated tubulin (AcT, yellow). (C) Overlay illustrating cell number and distribution by DNA staining (blue) and basal bodies stained with anti-γ-tubulin (γTub, cyan). Maximum intensity projection of 29 confocal sections. Scale bar A-C, 10 μm. The central canal lumen is outlined with a white dashed line (cc) in panel A.

**Figure S2.**
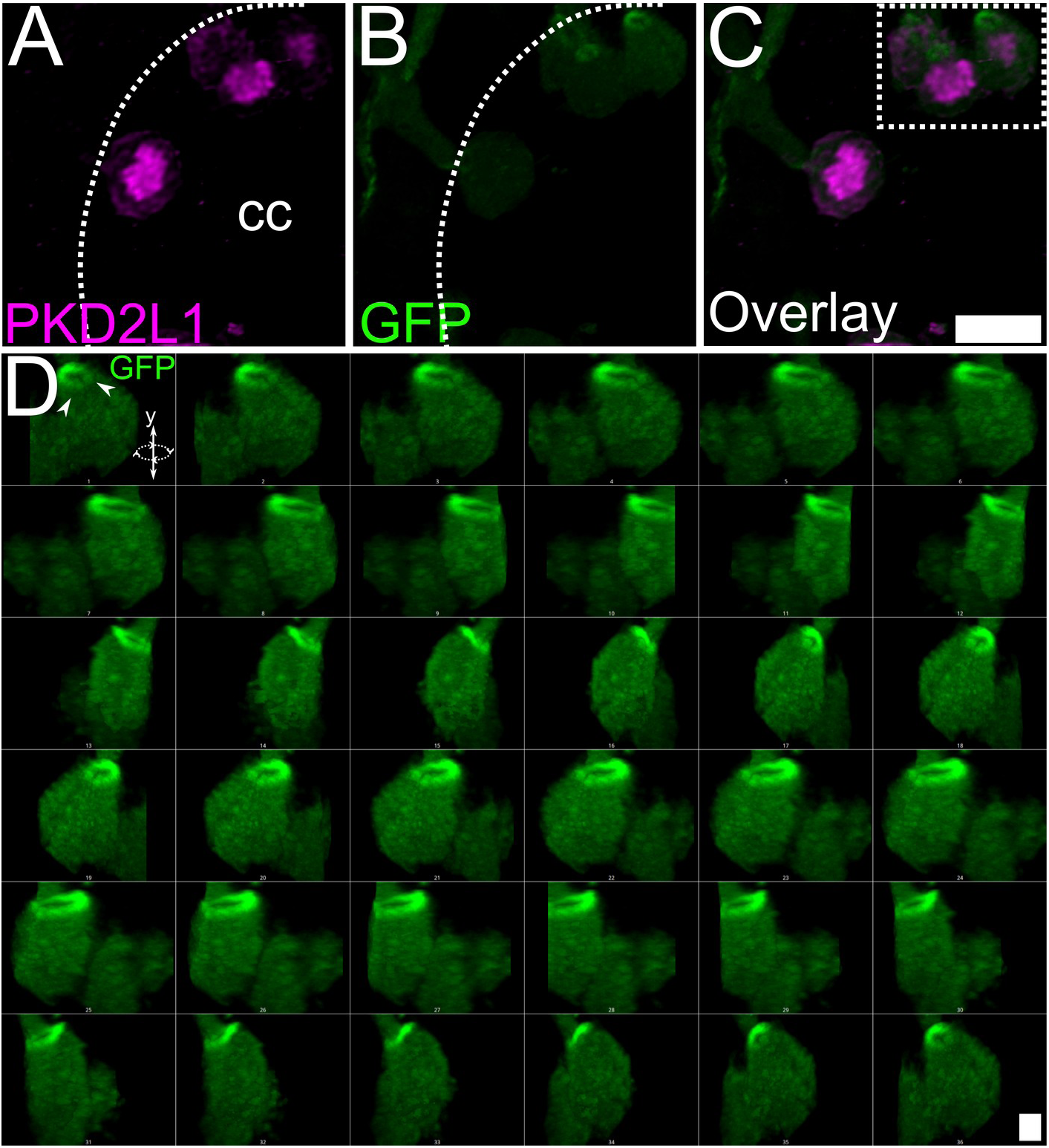
The ApPr neck forms a constricted domain enriched in cytosolic content, related to Figure 2. (A) PKD2L1 immunostaining reveals CSF-cN ApPrs protruding into the cc. (B) *GATA3^eGFP^*mice reveal a ring-shaped accumulation of GFP at the base CSF-cN ApPrs (arrowheads). (C) Overlay of A and B, highlighting the region reconstructed in D (white rectangle). (D) A 360° rotation around the y-axis of a 3D projection of a CSF-cN ApPr from a *GATA3^eGFP^* mouse, created from 45 deconvolved confocal planes (130 nm spacing). Rotation along the y-axis highlights the accumulation of the cytosolic GFP reporter at the constricted neck region (arrowheads). Scale bar A-C, 5 µm; D, 1 µm. The central canal lumen is outlined with a white dashed line (cc) in panels A and B.

**Figure S3.**
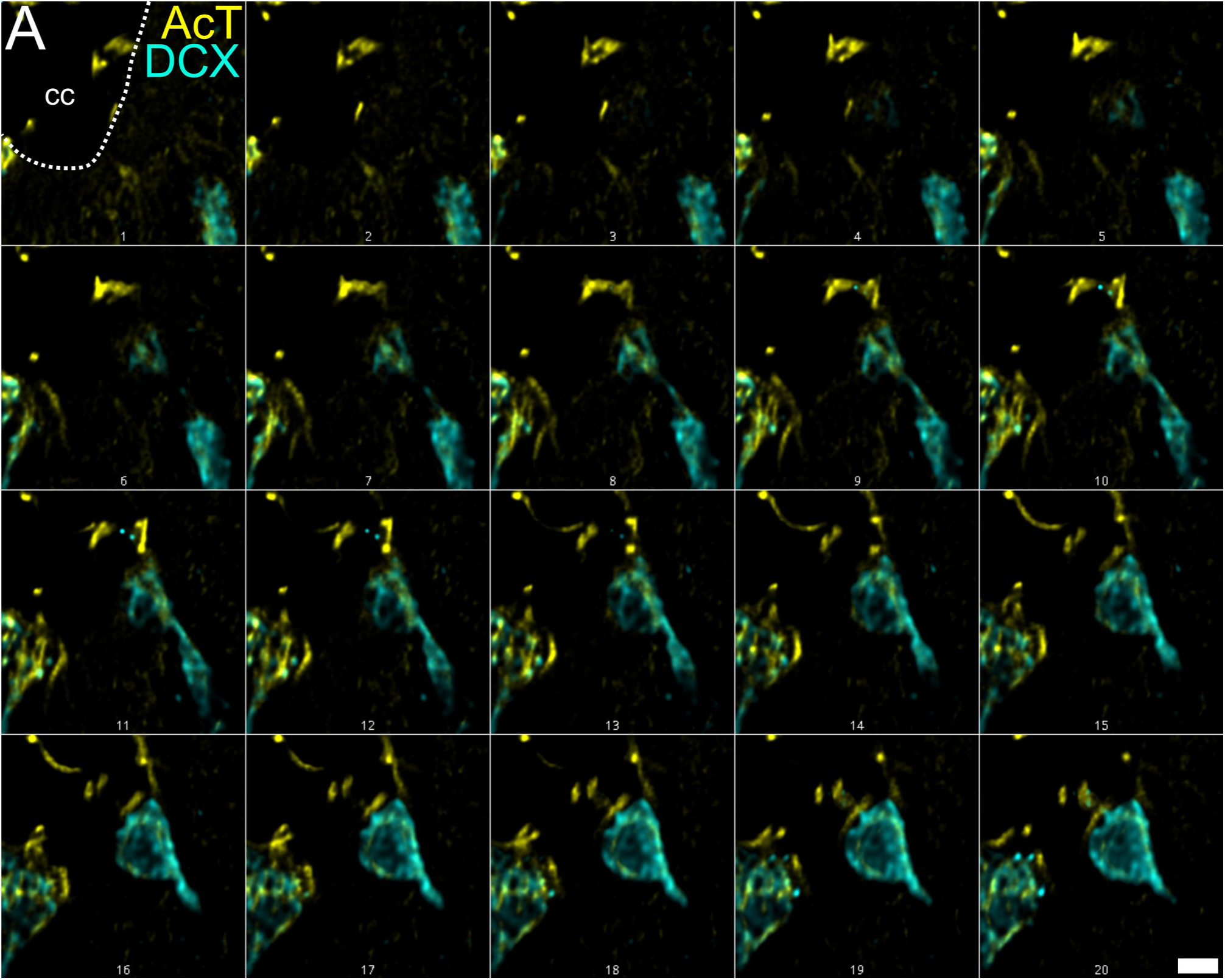
Doublecortin stabilizes a non-ciliary microtubule network in the ApPr, related to Figure 3. (A) High-resolution single confocal planes (deconvolved, 130 nm spacing) of mouse spinal cord showing the distribution of doublecortin (DCX, cyan) and acetylated tubulin stable microtubules (AcT, yellow) in CSF-cN apical processes (ApPrs). Scale bar, 2 µm. The central canal lumen is outlined with a white dashed line (cc) in panel A.

**Figure S4.**
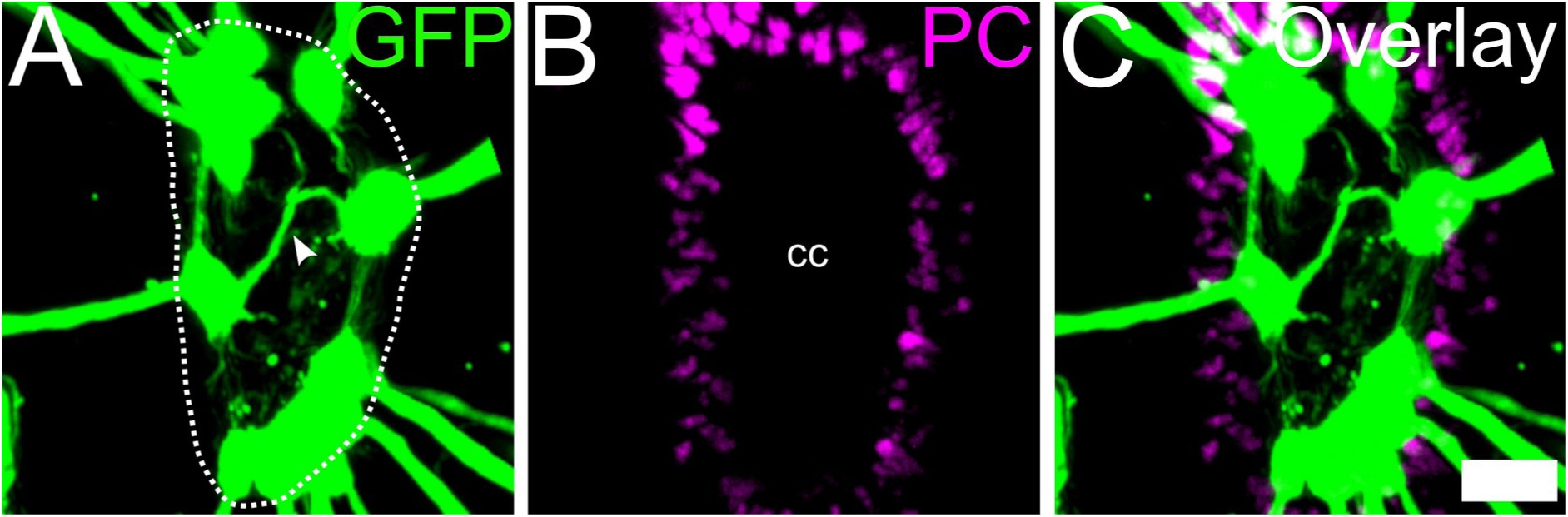
ApPrs emit interconnecting protrusions and exclude basal bodies, related to Figures 3 and 4. (A) A maximum intensity projection (65 z-planes, 500 nm spacing) of CSF-cN ApPrs (green) from a *GATA3^eGFP^* mouse shows fine protrusions (arrowhead) that appear to interconnect different ApPrs. (B) Staining for pericentrin (PC, magenta), a marker of basal bodies, reveals their presence in the surrounding tissue. (C) Overlay demonstrates that basal bodies are excluded from the GFP^+^ ApPrs, confirming their non-ciliated nature. Single confocal planes and a 3D reconstruction are shown in Supplementary Videos 3 and 4, respectively. Scale bar, 5 µm. The central canal lumen is outlined with a white dashed line (cc) in panel A.

