## Supplemental Video captions for "A filopodia-based dendritic mechanosensory compartment in CSF-contacting neurons"

Supplemental Information

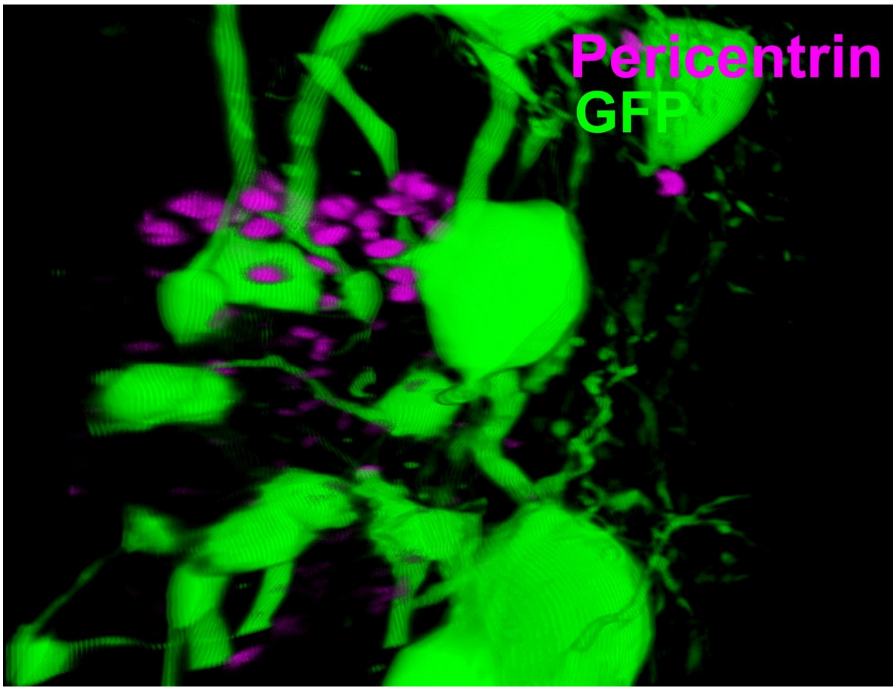


**Video S1. Apical Processes of CSF-cNs Are Cilia-Free and Lack Pericentrin-Positive Basal Bodies, Related to Figure 1 and S4.** Three-dimensional projection of a GFP-expressing spinal CSF-cN (green). Pericentrin immunostaining (magenta) labels basal bodies, which are absent from the GFP$^{+}$ apical process (ApPr). This shows that the basal bodies belong to adjacent ependymal cells.

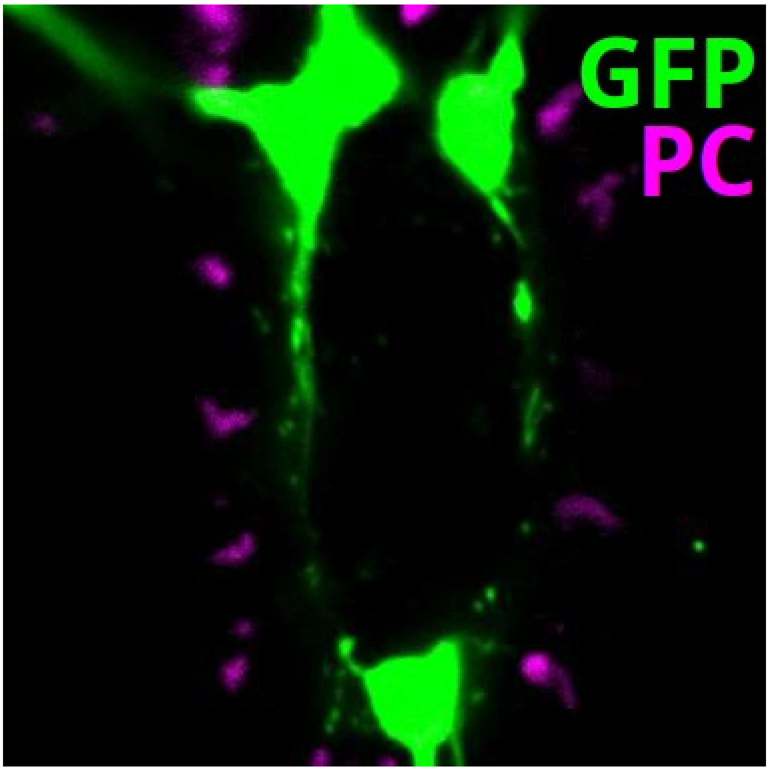


**Video S2. Confocal Z-Series Showing Exclusion of Basal Bodies from CSF-cN Apical Processes, Related to Figure 1 and Video S1.** Sequential confocal planes (Z-stack) used to generate the 3D projection in Video S1. GFP labels a CSF-cN (green). Pericentrin immunostaining (magenta) shows basal bodies are located adjacent to, but not within, the GFP+ apical process.

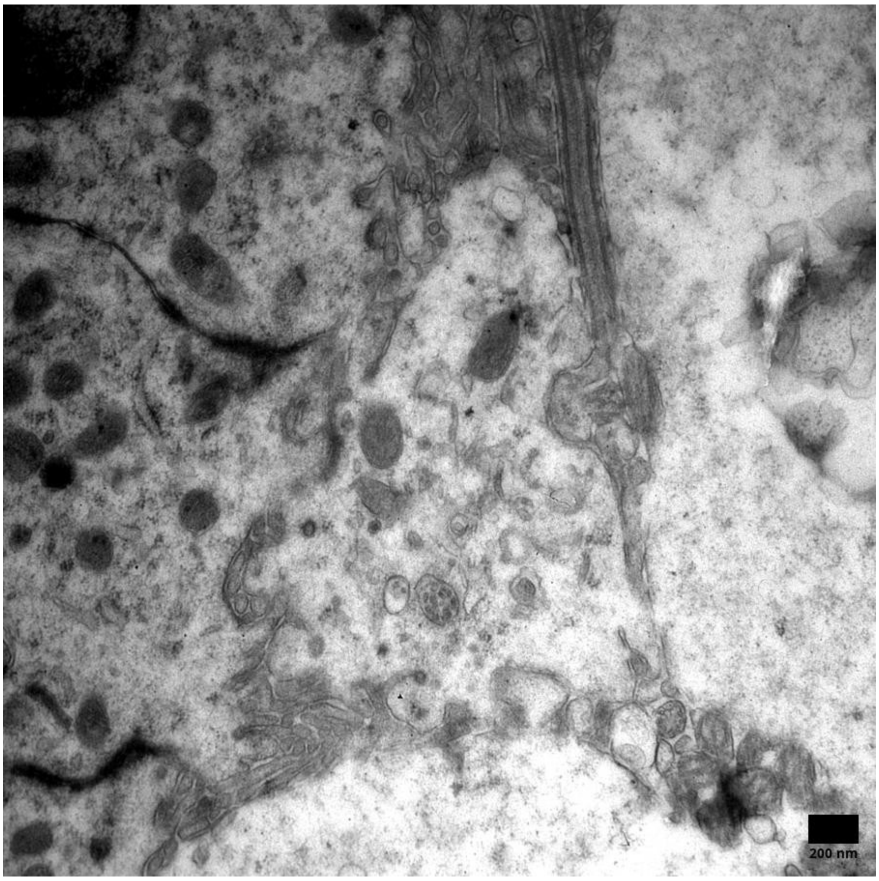


**Video S3. Ultrastructural Evidence that the CSF-cN Apical Process is a Cilia-Free Compartment, Related to Figure 1N.** Serial-section transmission electron microscopy (TEM) of a mouse spinal cerebrospinal fluid-contacting neuron (CSF-cN) apical process (ApPr) projecting into the central canal. This video steps through 26 consecutive ultrathin sections, revealing the detailed internal organization of the ApPr and its direct, non-ciliated interface with the cilia of adjacent ependymal cells. Scale bar, 200 nm.

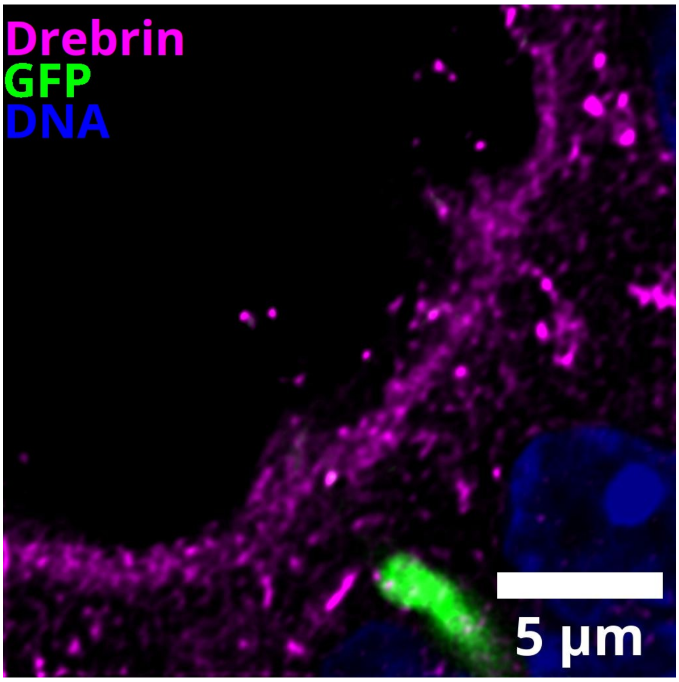


**Video S4. Drebrin Enrichment in the Actin-Cytoskeleton of CSF-cN Apical Processes, Related to Figure 3I and S4.** Deconvolved confocal Z-stack of the apical domain of GFP⁺ CSF-cNs (green), showing their processes extending into the central canal. Drebrin immunostaining (magenta) reveals strong enrichment along the apical process and its filopodia-like projections. DNA counterstain (blue) shows positions of ependymal cell nuclei.

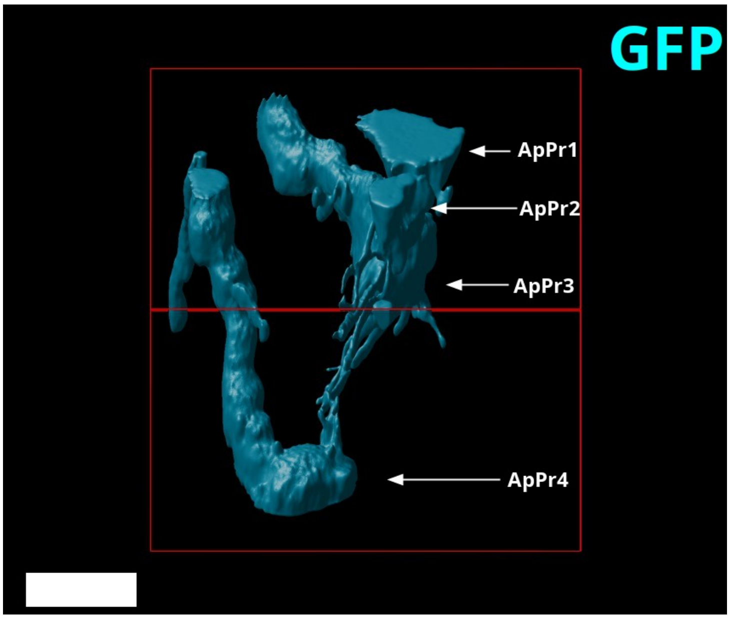


**Video S5. 3D Reconstruction Reveals Interconnections Between Neighboring CSF-cN Apical Processes, Related to Figures 3 and 4.** Three-dimensional reconstruction of the distal segments of four neighboring GFP⁺ CSF-cN apical processes (cyan; ApPr1–4). The rendering reveals fine terminal projections that appear to interconnect adjacent processes. Scale bar, 5 µm.

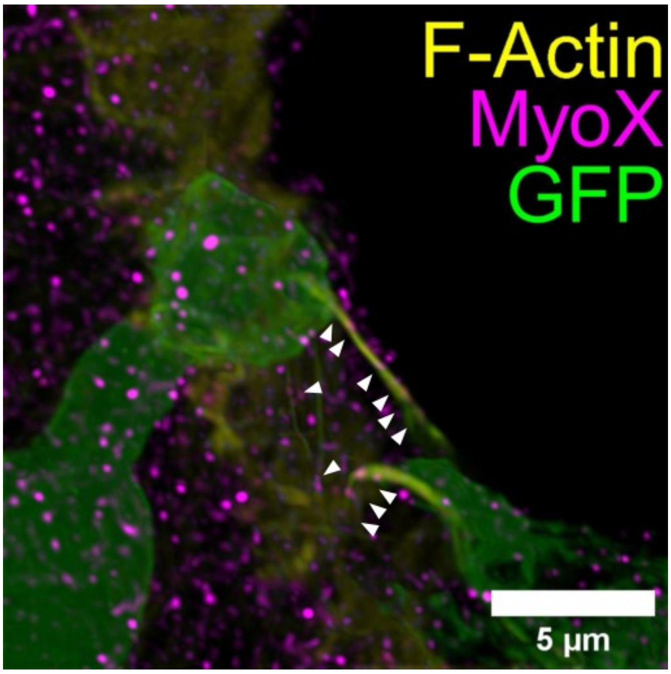


**Video S6. Apical Process major protrusions are enriched in F-Actin and Myosin-X, Related to Figure 4.** Deconvolved confocal Z-stack of the apical domain of GFP$^{+}$ CSF-cNs (green), showing their processes extending into the central canal. Immunostaining (magenta) reveals F-actin (yellow) is decorated with Myosin-X along the apical process and its finger-like protrusions (arrowheads). Scale bar, 5 µm.
